# Conscious perception of natural images is constrained by category-related visual features

**DOI:** 10.1101/509927

**Authors:** Daniel Lindh, Ilja G. Sligte, Sara Assecondi, Kimron L. Shapiro, Ian Charest

## Abstract

Conscious perception is crucial for adaptive behaviour yet access to consciousness varies for different types of objects. The visual system comprises regions with widely distributed category information and exemplar-level representations that cluster according to category. Does this categorical organisation in the brain provide insight into object-specific access to consciousness? We address this question using the Attentional Blink (AB) approach with visual objects as targets. We find large differences across categories in the AB then employ activation patterns extracted from a deep convolutional neural network (DCNN) to reveal that these differences depend on mid- to high-level, rather than low-level, visual features. We further show that these visual features can be used to explain variance in performance across trials. Taken together, our results suggest that the specific organisation of the higher-tier visual system underlies important functions relevant for conscious perception of differing natural images.

## Introduction

A long-standing question in cognitive neuroscience is how visual information is transformed from segregated low-level features to fully conscious and coherent representations. Prevailing object recognition models propose that rapid object identification is accomplished by extracting increasingly complex visual features at various stages/locations of the visual stream (DiCarlo, Yoccolan, & Rust, 2012; Felleman & Van Essen, 1991; Ungerleider & Haxby, 1994). Objects are first processed through a hierarchy of ventral visual areas where computations evolve from image feature detection, shape and part segmentation, before more invariant, semantic representations of the objects are established (Charest, Kievit, Schmitz, Deca, & Kriegeskorte, 2014; Cichy, Pantazis, & Oliva, 2014; Clarke & Tyler, 2014). The main question we address in the present report is how conscious access to objects differs as a function of category and visual features.

One particular categorical group that has been extensively studied is animacy, which show distinct processing pathways throughout the visual stream (e.g. Sha et al., 2015). Behavioural studies have shown that animate objects are more often consciously perceived in rapid serial visual presentations (RSVP) (Evans & Treisman, 2005; Guerrero & Calvillo, 2016; Hagen & Laeng, 2017), more quickly found in visual search (Jackson & Calvillo, 2013), elicit faster responses in discrimination tasks (Carlson, Ritchie, Kriegeskorte, Durvasula, & Ma, 2014; Ritchie, Tovar, & Carlson, 2015), and animate words are better retained in working memory (Nairne, VanArsdall, Pandeirada, Cogdill, & LeBreton, 2013). Aggregated, these findings point to a preferential visual processing of animate objects, most likely also reflected in the representational organisation of the visual stream (Carlson et al., 2014; Ritchie et al., 2015). However, the animate categorical division contains several sub-categories also known to cluster together, such as scenes in the parahippocampal place area (Epstein, Harris, Stanley, & Kanwisher, 1999), faces in the fusiform face area (Kanwisher, McDermott, & Chun, 1997) and body parts in the extrastriate body area (Downing, Jiang, Shuman, & Kanwisher, 2001; for review see Martin, 2007). It remains unclear how such sub-categories also might differ in visual processing. We address this question by testing differences across several categories, known to cluster together throughout the visual stream, in their propensity to conscious access in the Attentional Blink paradigm (AB; Raymond, Shapiro, & Arnell, 1992).

In the AB paradigm, two targets (T1 and T2) are embedded in a rapidly presented stimulus stream (RSVP). The frequently replicated finding is a reduced ability to report T2 when it is presented in a temporal window of 200-500 ms after a correctly identified T1. This effect disappears when subjects are asked to ignore T1 (Raymond et al., 1992), indicating that the fundamental explanation for this effect is attentional rather than perceptual. Most theoretical accounts of the AB suggest a two-stage information-processing model (Chun & Potter, 1995; for review see Dux, 2009). First, both targets are rapidly and automatically processed to a high-level representational stage. This is followed by a capacity-limited second stage, where the percept is transformed into a reportable state (i.e. working memory). Neural findings (Shapiro et al., 2007; Luck, Vogel, & Shapiro, 1996; Marois, Yi, & Chun, 2004; Sergent, Baillet, & Dehaene, 2005) have suggested that the AB is occurring at the second stage, after semantic processing of the object. This is in contrast to backwards masking, which is known to interrupt feedback loops in early processing (Fahrenfort, Scholte, & Lamme, 2009; Harris, Schwarzkopf, Song, Bahrami, & Rees, 2011; Kovács, Vogels, & Orban, 1995). Since feedback loops between visual areas are thought to be intact in the AB (Dehaene, Changeux, Naccache, Sackur, & Sergent, 2006), combined with a behavioural outcome that typically yields a significant number of both correct and incorrect trials, this paradigm is an ideal approach to investigate the bifurcation between conscious and unconscious visual processing.

One potential problem of studying categorical differences is that many categories share low-level scene statistics (Torralba & Oliva, 2003), which also are known to explain behaviour (Groen, Ghebreab, Lamme, & Scholte, 2012). Consequently, an issue that must be taken into account is how to control for low-level scene statistics in a neurally plausible way. We address this issue by using a Deep Convolutional Neural Network (DCNN; Krizhevsky, Sutskever, & Hinton, 2012) which is designed in a hierarchy encompassing feature representations of increasing complexity, similar to the visual system. Recent studies using DCNNs trained to classify a large corpus of natural images have revealed a significant correspondence between DCNN layers and the visual hierarchical organisation in the brain both using fMRI (Cichy, Khosla, Pantazis, Torralba, & Oliva, 2016; Eickenberg, Gramfort, Varoquaux, & Thirion, 2017; Güçlü & van Gerven, 2014; Khaligh-Razavi & Kriegeskorte, 2014), and MEG (Cichy et al., 2014; Greene & Hansen, 2018). This makes DCNNs attractive for modelling visual features rather than relying on manually labelling image features without knowing their relevant correspondence to the visual system.

**Figure 1:**
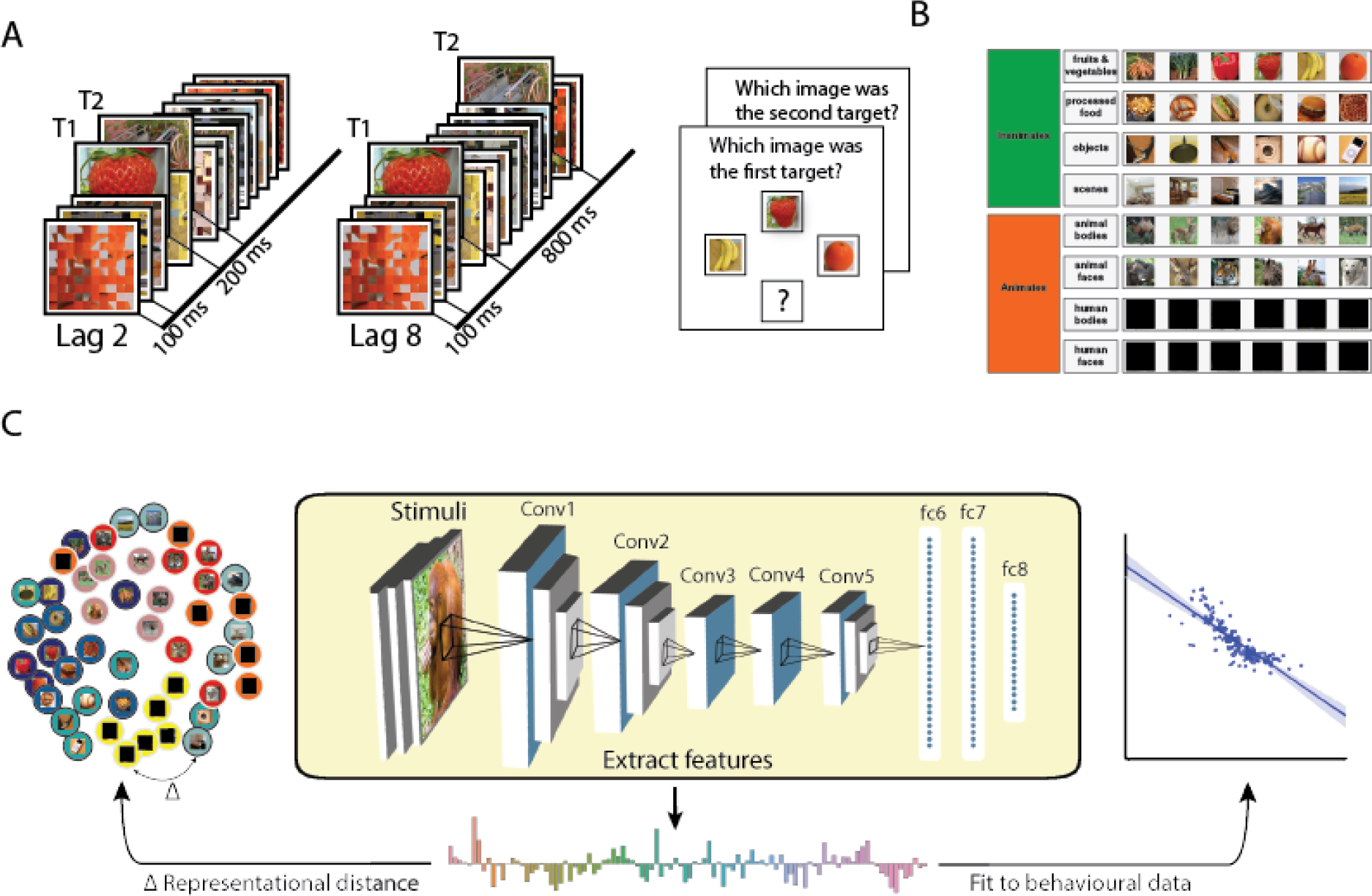
Modulating conscious access using the Attentional Blink Paradigm. A) We presented a rapid serial visual presentation to participants, with two targets (T1 and T2) following each other within a stream of distractors. On the left, the second target (T2) is shown 200ms after the first target (T1), and on the right, 800ms after the T1. In every trial, participants had to detect and later recall both T1 and T2 targets. B) The visual stimuli shown as targets to participants, consisted of animate and inanimate objects including exemplars of fruits and vegetables, processed foods, objects, scenes, animal bodies and faces, and human bodies and faces (these are blacked out for privacy protection reasons). Images were presented in grayscale and only visible here in colour for presentation purposes. C) We used a deep convolutional neural network (DCNN; yellow insert; 5 convolutional layers and 3 fully connected layers) to model the stimulus representational geometries (left) and predict our participants’ behaviour (right). The visual stimuli were fed into the DCNN, providing a hierarchical representation for each image. These unit activations were then analysed layer-by-layer and used to predict behaviour.

The main question of the current study was if the organisation of the visual system promotes conscious access to certain objects more than others. A priori, we had two related hypotheses. First we hypothesised that categories will differ in their access to consciousness. Our second hypothesis was that variance in conscious access between image exemplars could be predicted using high-level features derived from the DCNN. These two predictions are consistent with our current understanding of the categorical organisation of the ventral visual stream (Charest et al., 2014; Clarke & Tyler, 2014; Huth, Nishimoto, Vu, & Gallant, 2012; Kriegeskorte, Mur, Ruff, & Kiani, 2008), the high resemblance in representational geometry between the brain and DCNNs (Cichy et al., 2016; Eickenberg et al., 2017; Greene & Hansen, 2018; Güçlü & van Gerven, 2014), and theoretical models positing the AB as a disruption of late selection (Dux, 2009; Shapiro, Raymond, & Arnell, 1994). In addition, we explored whether trial-by-trial variance in performance was related to the similarity between the two targets in terms of visual features. We asked whether this relationship has any impact on conscious access, and if so, at what stage of processing do the two targets interact? To test this formally, we used a novel method coined *representational sampling*, where trials of the AB are constructed with stimuli selected according to their location in DCNN representational geometries. Our use of DCNNs overcomes earlier difficulties in characterising image features to better understand their contribution to conscious access in object recognition.

## Results

### Experiment 1

#### Differences in attentional blink magnitude as a function of category

First, we observed a significant AB effect with T2 performance differences between lags (Lag 2; accuracy M = 0.704, SD = 0.041, Lag 8; M = 0.847, SD = 0.129, t(18) = −6.427, p < 0.001, see Fig 2A). Our 48 images were derived from 8 different categories – fruits and vegetables, processed foods, objects, scenes, animal bodies, animal faces, human bodies and human faces. We first pooled the images according to animate and inanimate objects (see Table 2). Animate and inanimate objects have previously been shown to be differentially affected during the AB (Evans & Treisman, 2005; Guerrero & Calvillo, 2016; Hagen & Laeng, 2017). Similarly here, a repeated measures 2×2 ANOVA with lag and animacy as factors showed a main effect of lag (F(1,18) = 34.09, p < 0.001, *η*^2^ = 0.654) and animacy (F(1,18) = 27.72, p < 0.001, *η*^2^ = 0. 606) as well as a significant interaction effect (F(1,18) = 45.63, p < 0.001, *η*^2^ = 0.606). Thus, in accordance with previous studies, the AB was less pronounced for animate images. For each sub-category individually, using a repeated measures ANOVA, we observed a main effect of T2-lag (F(1,18)=42.87, p < 0.001, *η*^2^ = 0.704) and category (F(7,126) = 45.49, p < 0.001, *η*^2^ = 0.716), along with an interaction between category and T2-lag (F(7, 126) = 23.99, p < 0.001, *η*^2^ = 0.571). Beyond the expected AB effect, the interaction effects reveal that different categories exhibit different attentional blink magnitudes (ABM; difference in performance between lag 8 and lag 2). Separate AB effects were tested by contrasting lag 8 and lag 2 performance within each category using a two-tailed t-test (Fig 2C) – Fruits and Vegetables ((18) = 6.912, p < .001), Processed foods (t(18) = 6.748, p < .001), Objects (t(18) = 3.003, p = .004), Scenes (t(18) = 8.073, p < .001), Animal bodies (t(18) = 5.259, p < .001), Animal faces (t(18) = 2.712, p =.007), Human bodies (t(18) = 1.162, p = .13), Human faces (t(18) = 2.632, p = 0.008).

**Table 1:** Mean and SDs for T2 accuracy for each category.

| T2\|T1 |  |  |  |  |
| --- | --- | --- | --- | --- |
| Lag | Category | Mean | SD | N |
| Lag 2 | Fruits Vegetables | 0.651 | 0.199 | 19.000 |
|  | Processed Foods | 0.595 | 0.214 | 19.000 |
|  | Objects | 0.806 | 0.173 | 19.000 |
|  | Scenes | 0.406 | 0.234 | 19.000 |
|  | Animal bodies | 0.642 | 0.232 | 19.000 |
|  | Animals faces | 0.782 | 0.197 | 19.000 |
|  | Human bodies | 0.859 | 0.179 | 19.000 |
|  | Human faces | 0.886 | 0.133 | 19.000 |
| Lag 8 | Fruits Vegetables | 0.867 | 0.139 | 19.000 |
|  | Processed Foods | 0.853 | 0.110 | 19.000 |
|  | Objects | 0.893 | 0.079 | 19.000 |
|  | Scenes | 0.695 | 0.237 | 19.000 |
|  | Animal bodies | 0.822 | 0.159 | 19.000 |
|  | Animals faces | 0.858 | 0.134 | 19.000 |
|  | Human bodies | 0.879 | 0.153 | 19.000 |
|  | Human faces | 0.927 | 0.085 | 19.000 |

**Table 2:** Mean and SDs for T2 accuracy for animacy.

| T2\|T1 |  |  |  |  |
| --- | --- | --- | --- | --- |
| Lag | Animacy | Mean | SD | N |
| Lag 2 | Animate | 0.792 | 0.171 | 19.000 |
|  | Inanimate | 0.683 | 0.185 | 19.000 |
| Lag 8 | Animate | 0.872 | 0.126 | 19.000 |
|  | Inanimate | 0.871 | 0.097 | 19.000 |

**Figure 2:**
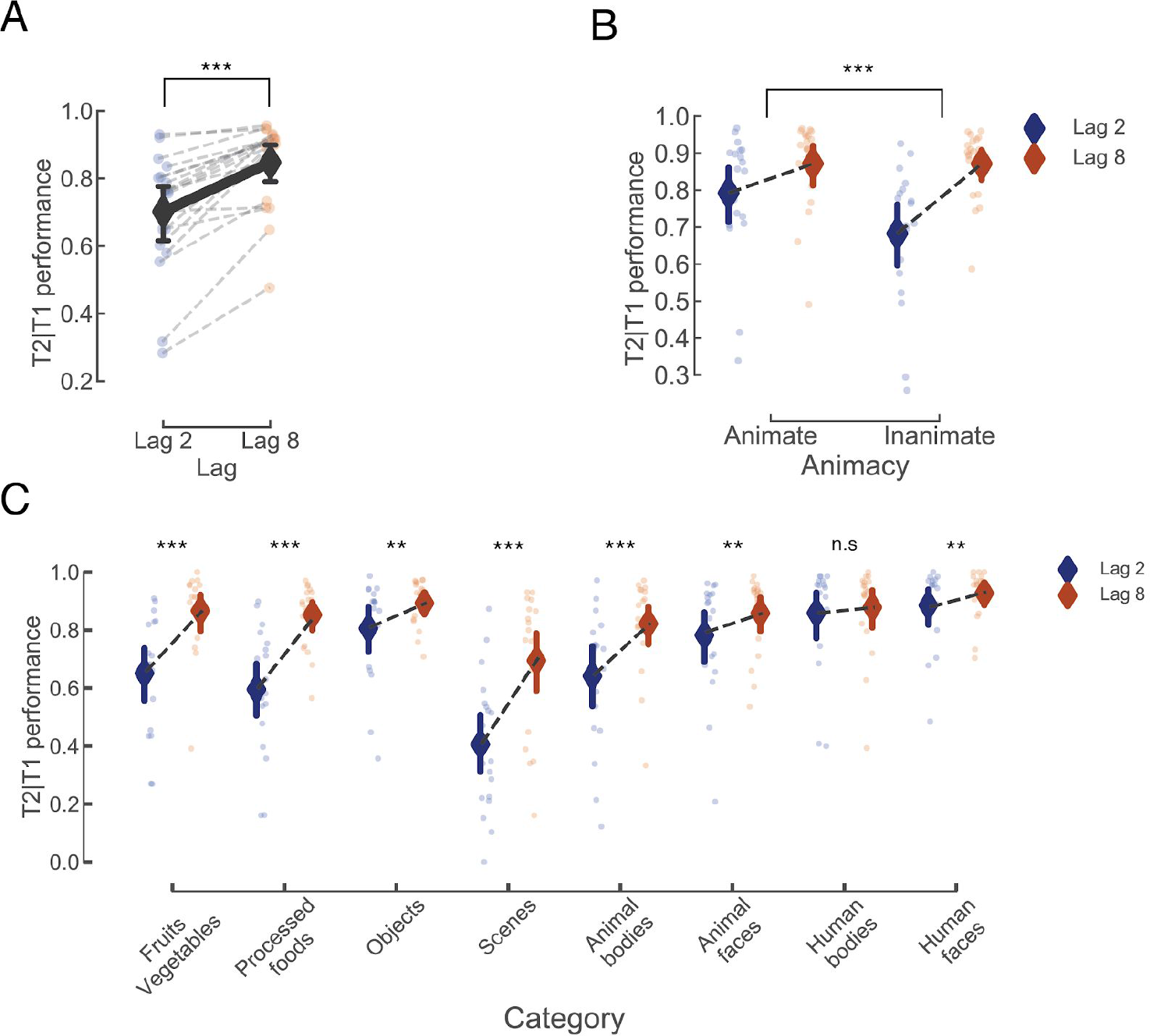
Animate objects elicit weaker attentional blink. A) The accuracy in detecting the second target conditional on having detected the first target for lag 2 and lag 8. Individual dots represent the mean performance for each subject, bold dots represent the mean performance across subjects, and error bars indicate 95% confidence interval around the mean in all plots. B) Performance plotted separately for animate and inanimate T2 targets. Attentional Blink Magnitude (ABM) is defined as the difference in performance between lag 8 and lag 2. Asterisk indicate significant difference in ABM between animate and inanimate. C) T2 performance for each category separately. Asterisks indicate p-values significant difference in ABM from zero. * = p < 0.05, ** = p < 0.01, *** = p < 0.001.

#### High-level image features within the DCNNs explain the majority of ABM variance between images and across trials

For each image we extracted unit activations from all the layers throughout an AlexNet DCNN (see methods). For the convolutional layers, we averaged over the spatial domain to obtain feature activations. It is important to note that this DCNN was trained on classifying objects into categories from a different set of images than those presented in our experiment, and at no point was trained on the AB. The feature activations were used in a cross-validated linear regression model aimed at predicting each image’s ABM. For significance testing we permuted the labels 3000 times for each subject and predicted the ABM based on the shuffled labels. We then averaged the prediction for each image across permutations and calculated the mean absolute error (MAE) for the image. We calculated the p-values by comparing this to the MAE of the non-permuted mean prediction from each layer across participants. After Bonferroni correction of the p-values, we were able to predict the ABM using features derived from layer conv4 (MAE = 0.07, pbonf = 0.008), conv5 (MAE = 0.061, pbonf < 0.001), fc7 (MAE = 0.069, pbonf < 0.001), and fc8 (MAE = 0.071, pbonf = 0.016). Testing the MAE of each participant per layer, we found that features from layer 7 is best suited for explaining the difference of AB effect across images (Fig 3B).

**Figure 3:**
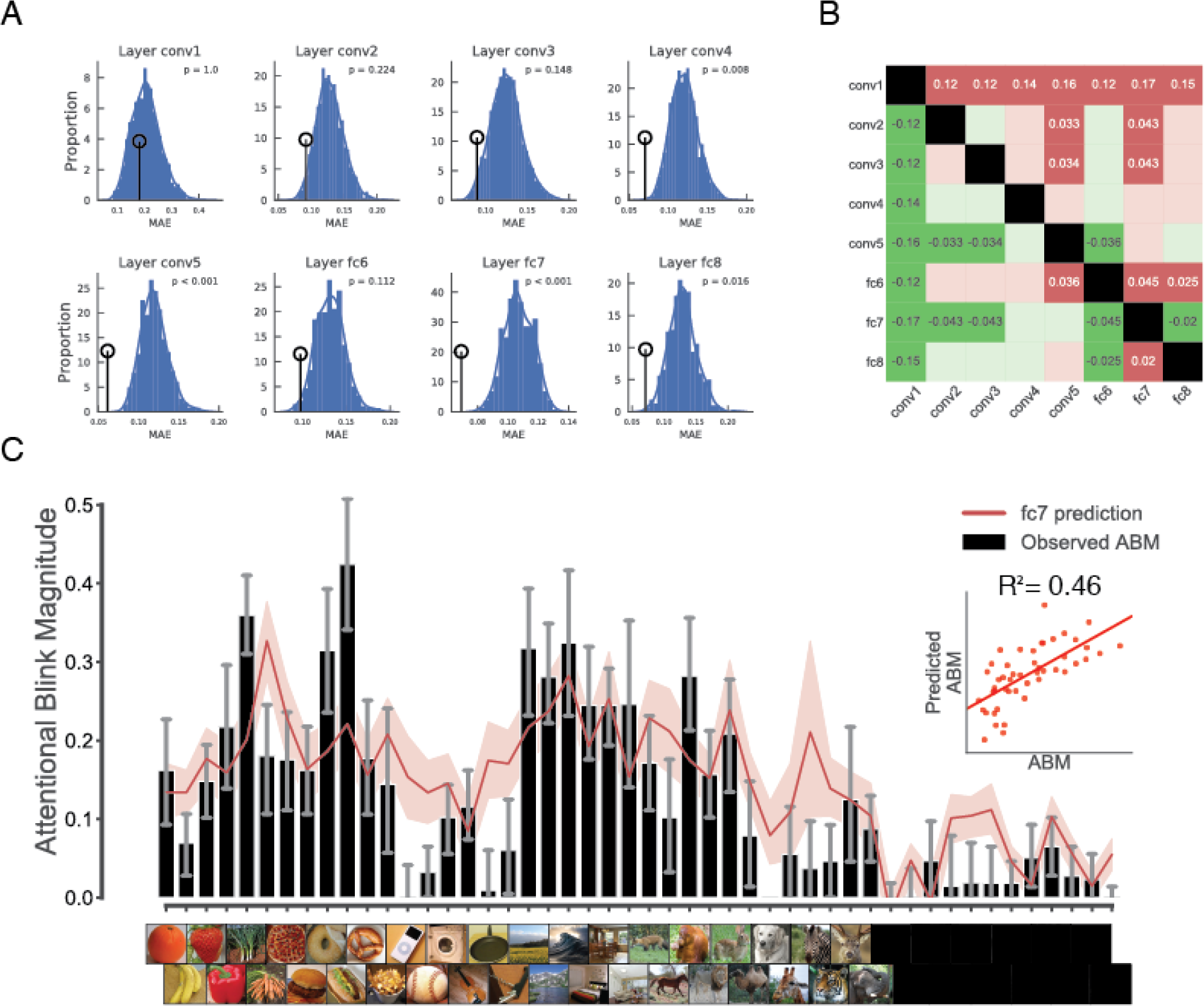
DCNN activation units predict attentional blink magnitude. A) Permutation test distributions. Histograms show the mean absolute error (MAE) after averaging the predictions across participants with randomised image labels. Circles point to the observed MAE. P-values, *Bonferroni corrected for multiple comparisons*, are denoted in each plot. B) Layer by layer comparisons of MAE. Comparisons are done row-wise, where green indicates a lower MAE, or better fit, in comparison to the corresponding column. Only significant (Bonferroni corrected) comparisons are denoted with mean difference in MAE between comparisons. C) ABM per image. Black bars indicate the observed ABM, red line is the average predicted ABM based on features from Layer fc7 (which outperformed all other layers, see panel B). Layer fc7 explained 46% of the variance observed. The insert panel shows the average predicted ABM on the y axis, and the average observed ABM per image, on the x axis.

#### Shared image features between targets predicts performance

In addition to predicting the ABM for each image, we sought to better understand the trial-by-trial differences in the AB. For each trial, we correlated the two targets (T1 and T2) based on their features (Pearson correlation, Fig 3B) to obtain a T1-T2 similarity measure based on each layer. We then correlated the T1-T2 similarity (from each layer) with the averaged T2 accuracy using Spearman correlation (Fig 4B). We tested the fisher transformed Spearman correlation from each subject against zero for each layer using a two-tailed t-test. Our results revealed that T1-T2 similarity significantly predicts T2 accuracy when based on layer conv2 (Spearman r, M = 0.22, SD = 0.21, t(17) = 4.41, p < 0.001), conv3 (M = 0.354, SD = 0.24, t(17) = 5.83, p < 0.001), conv4 (M = 0.15, SD = 0.212, t(17) = 2.93, p = 0.009), conv5 (M = 0.15, SD = 0.19, t(17) = 3.27, p = 0.004), fc6 (M = 0.18, SD = 0.17, t(17) = 2.77, p < 0.001), fc7 (M = 0.17, SD = 0.19, t(17) = 2.77, p = 0.013), fc8 (M = 0.29, SD = 0.16, t(17) = 7.32, p < 0.001). This suggests that the ongoing visual processing of T1 can lower the conscious access threshold for T2, if T2 shares visual features with T1. This was true for all layers except for layer 1.

**Figure 4:**
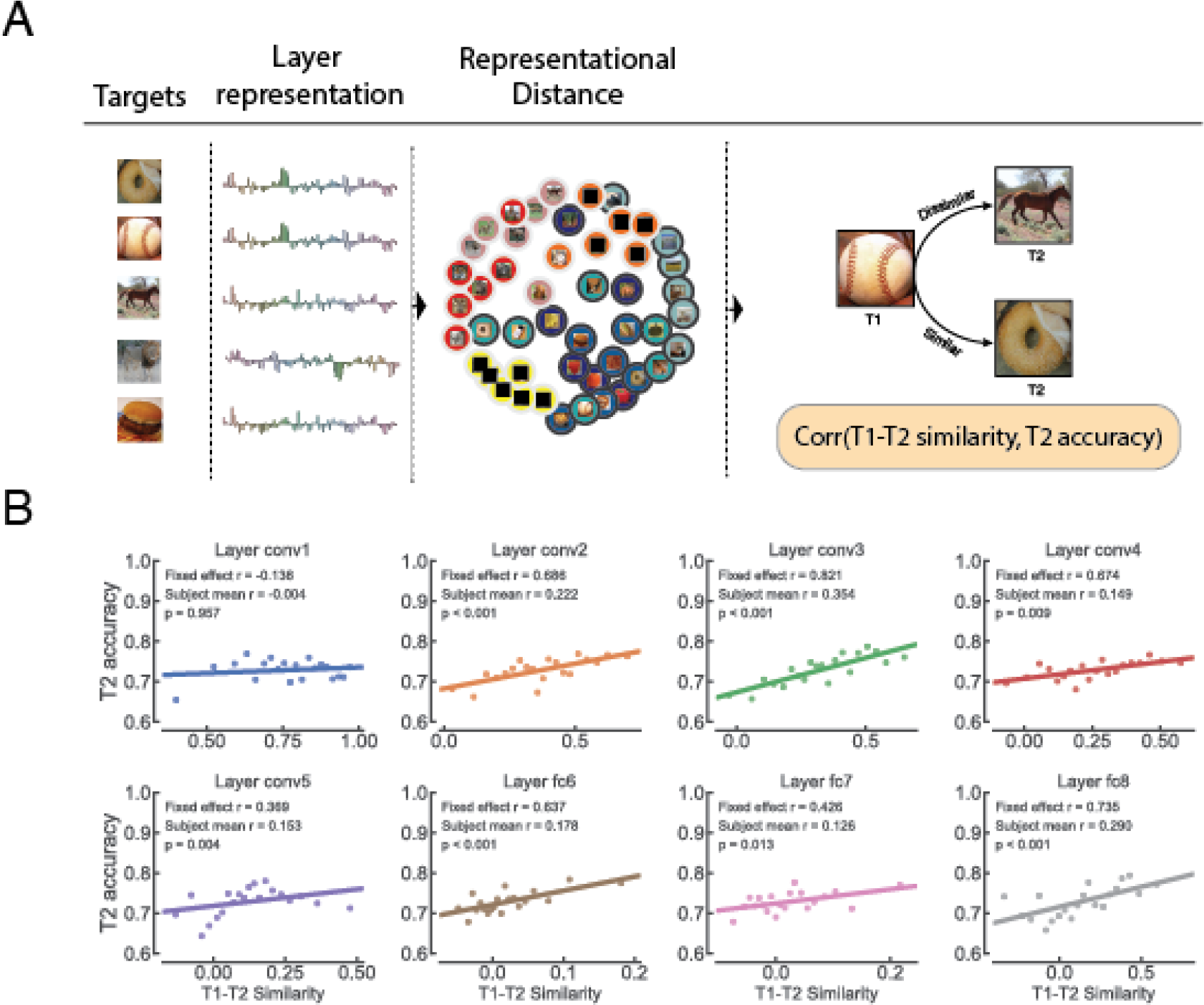
DCNN representational distance and target similarity explain trials of the AB. A) Depiction of analysis procedure. For each layer, DCNN representations are extracted for each image. These unit activations are then compared for all image pairs (Pearson correlation distance), to estimate the representational geometry. B) Correlation between lag-2 T2 performance and target-target similarity (based on 20 bins of similarities, averaged across participants) within each layer. Dots depict the binned similarities and lines depict regression line. The fixed effect r-value (based on the average bin values across subjects), the average r-value over subjects, and the p-value (obtained by testing the individual r-coefficients against zero) are denoted in each plot.

### Experiment 2

#### Constructing AB trials using representational sampling

The finding that T1-T2 similarity influences T2 accuracy prompted us to design a follow-up study. Because the results from the first experiment were correlational, we sought to investigate the effect of target-target similarity by manipulating the targets’ category and feature similarity. We developed a novel procedure called *representational sampling,* which first characterises a variety of stimulus response profiles, and samples a subset of stimuli tailored for our experiment. We used unit activations from layer 5 (see methods for rationale) of the DCNN as stimulus response profiles. We measured these unit activations on 250 images, derived from ImageNet (Russakovsky et al., 2015), to yield 16 images as our T2s; in turn chosen to represent four categorical groups equally (mammals, insects, vehicles, and furniture). For each image we then selected two T1s based on category (same or different) and similarity within layer 5 (similar or dissimilar), resulting in eight T1s per T2. This allowed us to examine the specific contribution of high-level feature similarity and category membership separately. We presented these four conditions to 24 new participants in an AB task similar to that of Experiment 1.

**Table 3:**
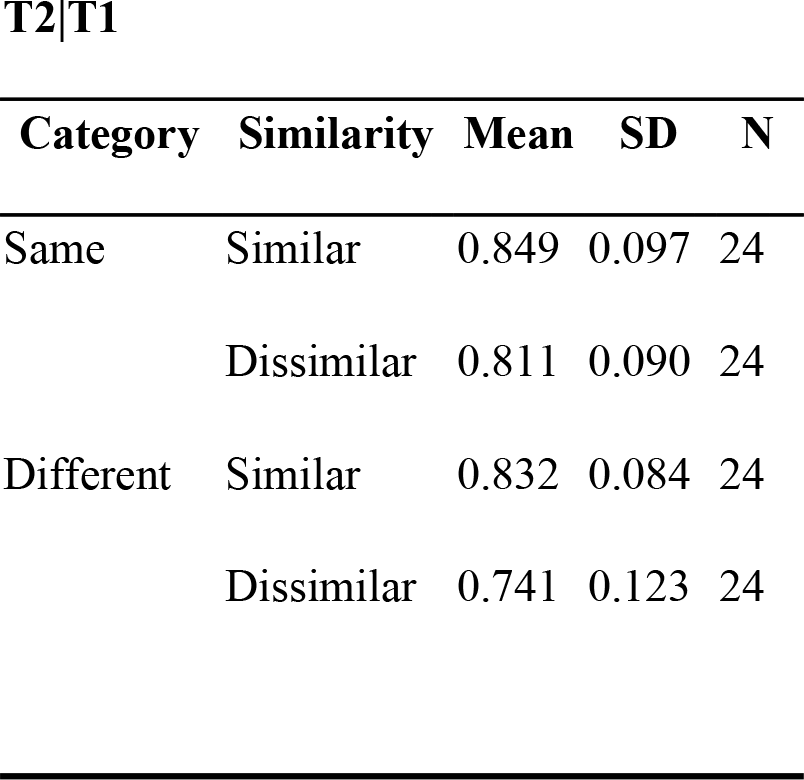
Mean and SDs for T2 accuracy in experiment 2.

Table 3 shows the group means of T2 performance for each of the four conditions. The probability of correctly reporting T2 was the highest when T1 came from the same category and had similar visual feature activation in layer 5 of the DCNN (M = 0.849, SD = 0.097). In contrast, the lowest probability of correctly reporting T2 was observed when T1 came from a different category and was dissimilar (M = 0.741, SD = 0.123). A 2×2 (Category by Similarity) repeated measure ANOVA showed a significant main effect for both category (F(1,23) = 20.68, p = <.001, *η*^2^ = 0.473) and similarity (F(1,23) = 45.468, p = <.001, *η*^2^ = 0.664), as well as an interaction effect (F(1,23) = 5.413, p = 0.029, *η*^2^ = 0.191). The larger effect size for the similarity factor indicates that visual features over semantic relevance determine behaviour.

**Figure 5:**
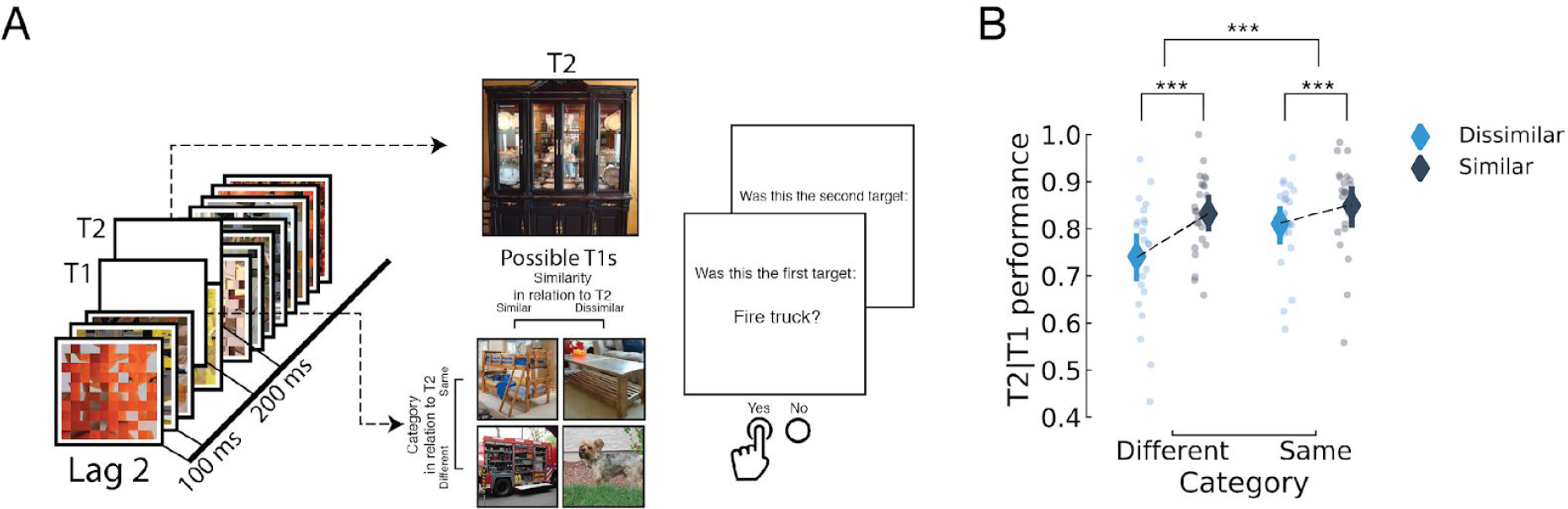
Target similarity between T1 and T2 explains T2 performance. A) Representational sampling was used to construct trials of experiment 2. Each of the sixteen T2s were either preceded by a T1 from the same/different category and similar/dissimilar in representational space within layer 5 of the DCNN. To ensure that participants did not use low-level statistics (such as colour) when reporting the targets, we switched the response menu to a semantic task. B) Behavioural results from experiment 2. Our results show that features similarity explain a significant portion of T2 performance. Individual dots correspond to individual subjects. Error bars indicate the 95% confidence interval. * = p < 0.05, ** = p < 0.01, *** = p < 0.001.

## Discussion

Here, we investigated the effect of category membership and image features on conscious access using natural images in the Attentional Blink paradigm (Fig 1A and B; Raymond et al., 1992). By testing images spanning several categories we first show a clear division in performance between animate and inanimate objects, where animate objects reveal a reduced AB caused by the processing of the T1 (Fig 2B), in line with previous reports (Evans & Treisman, 2005; Guerrero & Calvillo, 2016). We further show that this bias is not only expressed between this super-ordinate division, but also extends to various sub-categories. Using a DCNN as a model of the visual system, we show that mid- and high-level features in natural images (Fig 3) regulate the AB magnitude. In addition, we show that target-target similarity (Fig 4 and 5) interacts with target selection, providing a mechanistic explanation of the AB phenomenon and of conscious access in object recognition.

Previous studies have shown differences between categories in the AB, most extensively between animate and inanimate objects (Einhäuser, Koch, & Makeig, 2007; Evans & Treisman, 2005; Guerrero & Calvillo, 2016; Hagen & Laeng, 2017). The animacy bias in visual processing has been attributed to evolutionary relevance, as opposed to visual expertise, reflected in its importance for ancestral hunter-gatherer societies (The animate monitoring hypothesis; New, Cosmides, & Tooby, 2007). Evidence for this hypothesis comes from a wealth of behavioural studies showing that animate objects are more quickly and more often detected in different types of attentional tasks (e.g., Jackson & Calvillo, 2013; New et al., 2007). Likewise, animate and inanimate objects are distinctly represented throughout the ventral visual stream (Sha et al., 2015; Wen et al., 2017), which has been argued to be an evolutionary phenomenon and not contingent on visual experience (Mahon et al., 2009). In our current study, we find that the AB magnitude (ABM – performance difference between Lag-8 and Lag-2) is larger for inanimate objects, similar to Guerrero and Calvillo (2016). The finding by Guerrero and Calvillo has been contested by Hagen and Laeng (2017) who showed that animate objects are simply reported more often, but that the ABM is unaffected. Our results argue against the findings of Hagen and Laeng and, more importantly, reveal that differences in AB magnitude exist in a myriad of sub-categories. To the best of our knowledge, we are the first to examine a significant number of categories, which are known to cluster throughout the visual cortex. We show a high variance in the effect of the AB across categories (Fig 2C), implying that distinctive sub-categories have special privilege in the path to conscious access. It is important to note that by looking at the differences between Lag-8 and Lag-2, effectively baselining each image with its own Lag-8 performance, our results cannot be explained by differential effects of masking. Importantly, this implies a dissociation between attentional relevance and conscious access, since it would be reasonable to assume that attentional relevance would affect Lag-2 and Lag-8 equally.

The finding that the ABM varies across categories (Fig 2C) is hard to interpret without properly examining image features of different complexities. Many semantic categories share low-level statistics (Groen et al., 2012; Groen, Silson, & Baker, 2017; Torralba & Oliva, 2003) and, without delving further than categorical membership, one cannot disentangle at which level of processing the differences occur. The prediction of ABM across visual objects achieved by modelling DCNN unit activations from the mid to late layers explained a large proportion of AB variance across images (~46% of the variance in layer fc7, Fig 3C). This implies that the bottleneck produced by the AB is due to late visual processing and probably reflects the particular categorical organisations within higher-tier visual areas. This relationship between neural representation of images and behavioural outcomes is supported by recent work showing that the particular representational organisation in late visual areas predicts certain behavioural measures, such as reaction time (Carlson et al., 2014; Grootswagers, Cichy, & Carlson, 2018; Ritchie et al., 2015). This ‘conceptual’ approach to conscious access promotes a more fundamental view to how visual consciousness might operate by focusing on the organisation of the visual system rather than on top-down mechanisms.

Our experiments further enabled us to explore the importance of T1-T2 similarity. Only a handful of studies have investigated target-target similarity in the context of AB (e.g., Awh et al., 2004; Einhäuser et al., 2007; Evans & Treisman, 2005; Serences, Scolari, & Awh, 2009; Sy & Giesbrecht, 2009). In one of the earliest attempts to study target-target similarity and its effect on T2 performance, Awh et al. (2004) concluded that similarity between targets is detrimental to T2 reportability. This lead to the proposal of the multiple-resource channel hypothesis (MRCH; Awh et al., 2004). According to the MRCH, two targets (T1 and T2) can be processed in parallel, but only if their visual features are different enough to be processed through distinct feature channels. While a few following studies have corroborated this notion (Einhäuser et al., 2007; Serences et al., 2009; Sy & Giesbrecht, 2009), our study reveals that similarity is beneficial for performance. The difference in results might be explained by our way of defining similarity by image features, rather than categorical membership, and thus it is possible that our findings reflect a priming effect not found in the previous studies (however, see Evans & Treisman, 2005). One related finding by Nieuwenstein, Chun, van der Lubbe, and Hooge (2005) showed that a preceding distractor sharing visual features with the T2 reduced the AB. Because our distractors were scrambled images and not natural images, it is possible that we see a similar type of cueing effect, where the T1 cues the T2. The combined findings of all these studies highlight an unexplored aspect of AB, where the relationship between the targets might play a major role in explaining many AB phenomena. Further questions could be explored using a combination of brain measures to determine representational similarity within subjects, which might potentially also explain individual differences in performance.

In conclusion, we present compelling evidence that DCNNs can model the visual system to predict human behaviour. Specifically, we present findings that attribute differences in conscious access between image exemplars to difficulties in representational readouts of features in higher-tier visual areas. This visual feature-related bias is reflected in a stable functional organisation, where fined-grained category distinctions have a larger impact on conscious access than previously believed. Moreover, we point to a more dynamic way in which the context (i.e. the similarity between T1 and T2) biases the probability for a target to be consciously perceived. In summary, our findings suggest that object categories and visual features processed in higher level visual ventral stream constrain conscious perception of natural images.

## Methods

### Experiment 1

#### Experimental procedure

Twenty participants (19 females; age range: 19-22; mean = 20.1 ± 1.2) were recruited for the study. We excluded two participants due to incomplete data. One additional participant was excluded for the image-by-image analyses due to lack of trials where T2 was correct for one image after filtering for T1 correct. All participants provided and signed informed consent and were rewarded for their time via course credits or financial compensation (at the standard rate of £7/h). All participants had normal or corrected-to-normal vision, and no known history of neurological disorders. The Ethical Review Committee of the University of Birmingham approved the experiment.

#### Procedure

Participants viewed visual objects in a rapid serial visual presentation (RSVP), and were asked to detect two targets (T1 and T2) embedded into a stream of distractors (Fig 1A). Following the stream, a response menu was presented for T1, which included the T1 and two foils, and the participant had to identify the target with a button press. A similar response menu was then presented to identify the T2. The foils in the menu always belonged to the same category as the targets (Fig 1A, right panel).

#### Design and Stimuli

Participants were seated 60 cm away from a Stone monitor (60Hz refresh rate), and stimuli covered 5 degrees of visual angle centrally on a grey background. Stimulus presentation was achieved using the Psychtoolbox extension (version 3; Brainard, 1997) in MatLab 2016b (MathWorks Inc., Natick, USA). Stimuli consisted of 48 images, derived from eight different categories: fruits and vegetables, processed foods, objects, scenes, animal bodies, animal faces, human bodies, and human faces (Fig 1B). To generate the items used as distractors in the stream, each quadrant of the image was divided into 25 squares, which were then inverted, and randomly assigned a new position in their quadrant. This procedure has been described previously (Marois, Yi & Chun (2004)). Following a standard Attentional Blink (AB) paradigm (Raymond, Shapiro, & Arnell, 1992), each trial started with 300 ms of fixation, followed by a rapid serial visual presentation (RSVP) consisting of 19 images. Each image was presented for 16ms with a stimulus-onset asynchrony (SOA) of 100 ms (Fig 1A). Embedded into the stream of distractors, two non-scrambled targets (T1 and T2) were presented at two different lag conditions (Lag-2: 200ms and Lag-8 - 800ms). The T1 was always presented as item 5 in the stream, while T2 was either presented as item 7 (Lag 2) or item 13 (Lag 8). All 48 images were presented on an equal number of trials either as T1 or as T2, randomized within blocks with no trial having the T1 and T2 coming from the same superordinate category. Importantly, the same pair of T1 and T2 was always presented in both the Lag-2 and Lag-8 conditions, within the exact same stream of distractor masks in the RSVP trial. Participants had to press one out of three buttons to identify the correct target from the foils, or a fourth button when they missed the target. The two foils came from the same category as the target.

#### Deep Convolutional Neural Network (DCNN)

We employed a DCNN (AlexNet; Krizhevsky et al., 2012; see fig 1C), implemented through Python and Caffe (Jia et al., 2014), as a model of the visual cortex for extracting hierarchical visual features from our stimuli. We chose AlexNet due to its relative simplicity, compared to more recent DCNNs, and its well-studied relation to the human visual system (Cichy, Khosla, Pantazis, Torralba, & Oliva, 2016; Khalig-Razavi & Kriegeskorte, 2014; Wen, Shi, Zhang, Lu, & Liu, 2016). It consists of eight layers of artificial neurons stacked into a hierarchical architecture, where preceding layers feed-forward information to the next layer (Fig 1B). The first five layers are convolutional layers, whereas the last three are fully connected layers. While the fully connected layers (fc6, fc7, and fc8) consist of one dimensional arrays (sizes of 4096, 4096, and 1000 units respectively), the convolutional layers have the dimensionalities of: layer 1 (conv1) - 96×55×55 (96 features, over 55 x 55 “retinotopical” units), layer 2 (conv2) – 256×27×27, layer 3 (conv3) – 384×13×13, layer 4 (conv4) – 384×13×13, and layer 5 (conv5) – 256×13×13. For all analyses we averaged the values in the convolutional layers for each image over the spatial dimension, leaving them with the vector length of 96, 256, 384, 384, and 256 respectively. This network was pre-trained on 1.3 million hand-labelled, natural images (ImageNet; Russakovsky et al., 2015) for classification into 1000 different categories (available at http://caffe.berkeleyvision.org/model_zoo.html), reaching near-human performance on image classification (Krizhevsky et al., 2012). Our test set of 48 images were analysed through the network, and we used the last processing stage of each layer as model output for further analyses.

#### Analyses of behaviour and image features

For each image, we calculated the mean T2 accuracy at both Lag-2 and Lag-8 across subjects. We then computed Attentional Blink Magnitudes (ABM) by subtracting Lag 2 mean accuracies from Lag 8 mean accuracies. ABM then becomes a measure of how much the AB time window affects the recall of each image separately. In the interest of quantifying image features, within our DCNN, we extracted unit (“neuron”) activation patterns for each image from all the layers. For the first five convolutional layers, we averaged the activation over the spatial dimension. These activation patterns were incorporated into a multivariate linear regression model (Sci-kit learn; Buitinck et al., 2013.), with the activation patterns from each layer as features in the model to predict each image’s ABM within subjects. The prediction pipeline followed a “leave-one-out” procedure – where, based on the training data, the features were first standardized to unit variance and feature selection was done by removing low-variance features.

#### Target-target similarity

We further tested the effect of target-target similarity on conscious access. Here, we go beyond using predetermined categories as a proxy for feature similarity and examined the representational distance between images within a given layer of the DCNN. The T2-performance to T1/T2-similarity correlation was implemented using the following steps for each DCNN layer. (A) For each trial, we correlated the T1 and T2 images within the given DCNN layer. (B) We binned all trials into 20 bins based on the correlation between the targets on any given trial, ensuring an equal number of trials in each bin. (C) We correlated the target-target similarity with T2 performance using Spearman’s r.

### Experiment 2

#### Participants

We recruited 24 participants (Age - M= 19.38, SD = 0.95, females = 19, males = 5).with normal, or corrected-to-normal, vision. All participants provided and signed informed consent and were rewarded for their time via course credits or financial compensation (at the standard rate of £7/h). The experiment was approved by the ethics committee at the University of Birmingham.

#### Procedure and stimuli

Unless stated otherwise, all procedure and visual presentations were identical to Experiment 1 (see fig 5A). Sixteen images, a subset of 250 labelled and processed images from the ImageNet database (Russakovsky et al., 2015), were selected as T2s. The T2s derived from four different categories (mammals, insects, vehicles, and furniture), and each category was uniformly represented in the T2 selections. Similarity between images was determined by their Pearson correlation coefficient within layer 5 of the DCNN. The layer 5 was chosen because it was a high-performing layer in the first study and to still maintain the retinotopical information for an additional analyses not used in this study. To model the layer-wise unit activations for this new set of images, we used the same pre-trained network (AlexNet, Krizhevsky et al. 2012) as in Experiment 1. For each T2, we selected two similar and two dissimilar images from the same category and any of the other categories as T1. This resulted in eight potential T1s for each T2 in a 2-by-2 factorial design (Similarity X Category) (Fig 5A). The T1 was always placed on position 11, and T2 on position 13 (in a RSVP of 19 items for each trial). Each block consisted of a presentation of each T2 paired with every possible T1, for a total of 128 trials per block divided into 4 runs (32 trials per run). Each participant completed 2 blocks for a total number of 256 trials per session (64 trials per condition).

## Acknowledgements

This work was supported by an European Research Council (ERC) Starting Grant ERC-2017-StG 759432 (to I.C.). We would like to thank Sara Binks and Alfie Brown for their help in collecting the behavioural data from experiments 1 and 2 (respectively), and Jasper van den Bosch for comments on the manuscript.

Appendix A. Supplementary data and code

Supplementary data and code associated with this article can be found, in the online version, at https://github.com/Charestlab/abdcnn/.

